# An *In-Vitro* Standardized Protocol for Preparing Smoke Extract Media from Cigarette, Electronic Cigarette and Waterpipe

**DOI:** 10.1101/2024.08.07.606957

**Authors:** Amel Al-Hashimi, Jagrut Shah, Roger Carpenter, Winston Morgan, Mohammed Meah, Prashant J. Ruchaya

**Author notes:** **Corresponding author Dr. Prashant J. Ruchaya**, University of East London, Water Lane, London, England, E15 4. joint first co-authors (Amel Al-Hashimi, Jagrut Shah).

## Abstract

Cigarette smoke exposure is a major risk factor for various diseases, and in vitro studies using smoke medium (SM) are crucial for elucidating underlying mechanisms. However, the lack of standardized protocols for SM preparation has led to inconsistencies in experimental results. This paper presents an optimized protocol for reproducible generation of SM from conventional cigarette smoke extract (CSE), electronic cigarette smoke (ECS), and waterpipe vapour (WPV). The protocol involves:

A custom-built apparatus for smoke capture in a cell culture medium
Specific parameters mimicking human smoking topography
Rigorous aseptic techniques and safety measures

Extensive details are provided for material preparation, apparatus setup, smoke generation procedures, and post-processing steps. This standardized method enables reliable in vitro exposure models, easing comparative studies across different tobacco products. To prove protocol consistency, we present optical. Additionally, we highlight the protocol’s utility by figuring out the SM quality through analysing O.D. data and morphological changes, cell viability assays, ELISA and gene expression analyses i.e., RNA sequencing through several *in-vitro* models, i.e., Human Umbilical Vein Endothelial Cells, (HUVECs, **Merck)**, human leukaemia monocytic cell line (THP-1, Sigma) and induced pluripotent stem cells differentiated to cardiomyocytes (iPSC-CMs, **Axol)** respectively. This optimized protocol addresses variability in SM preparation, enabling consistent investigations into smoke-induced pathologies and potential therapeutic interventions.

**Graphical abstract:** 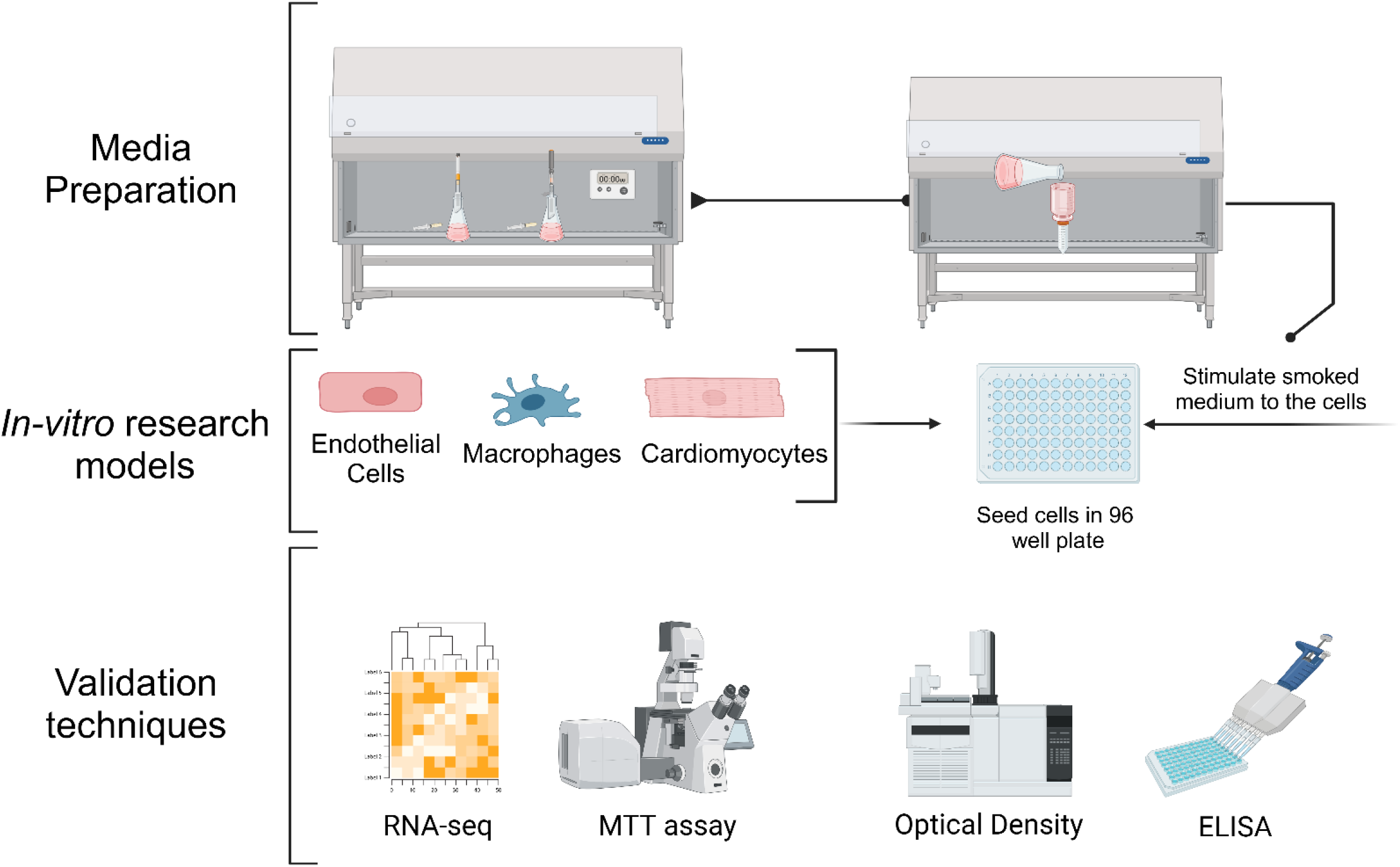

**Specifications table:** 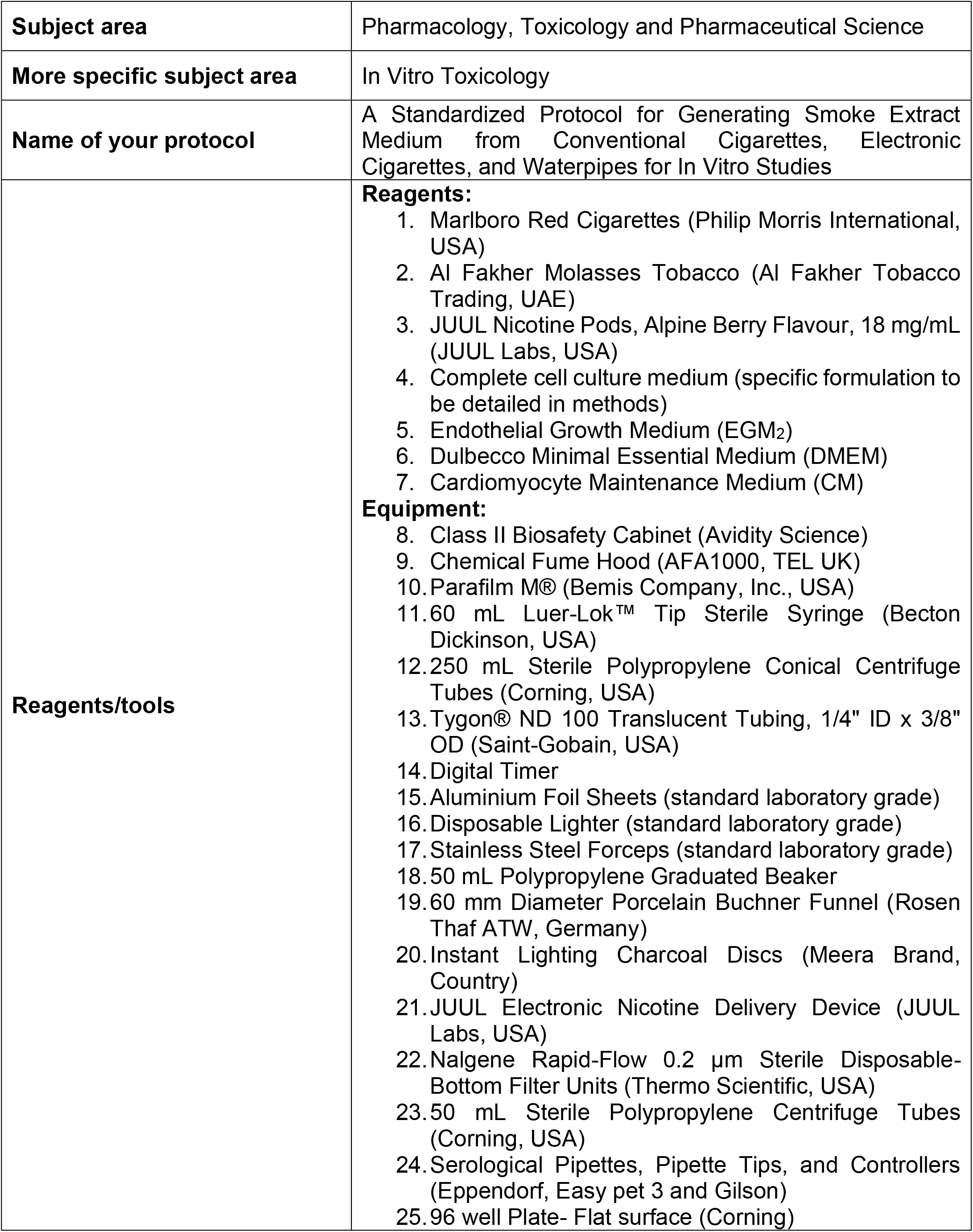

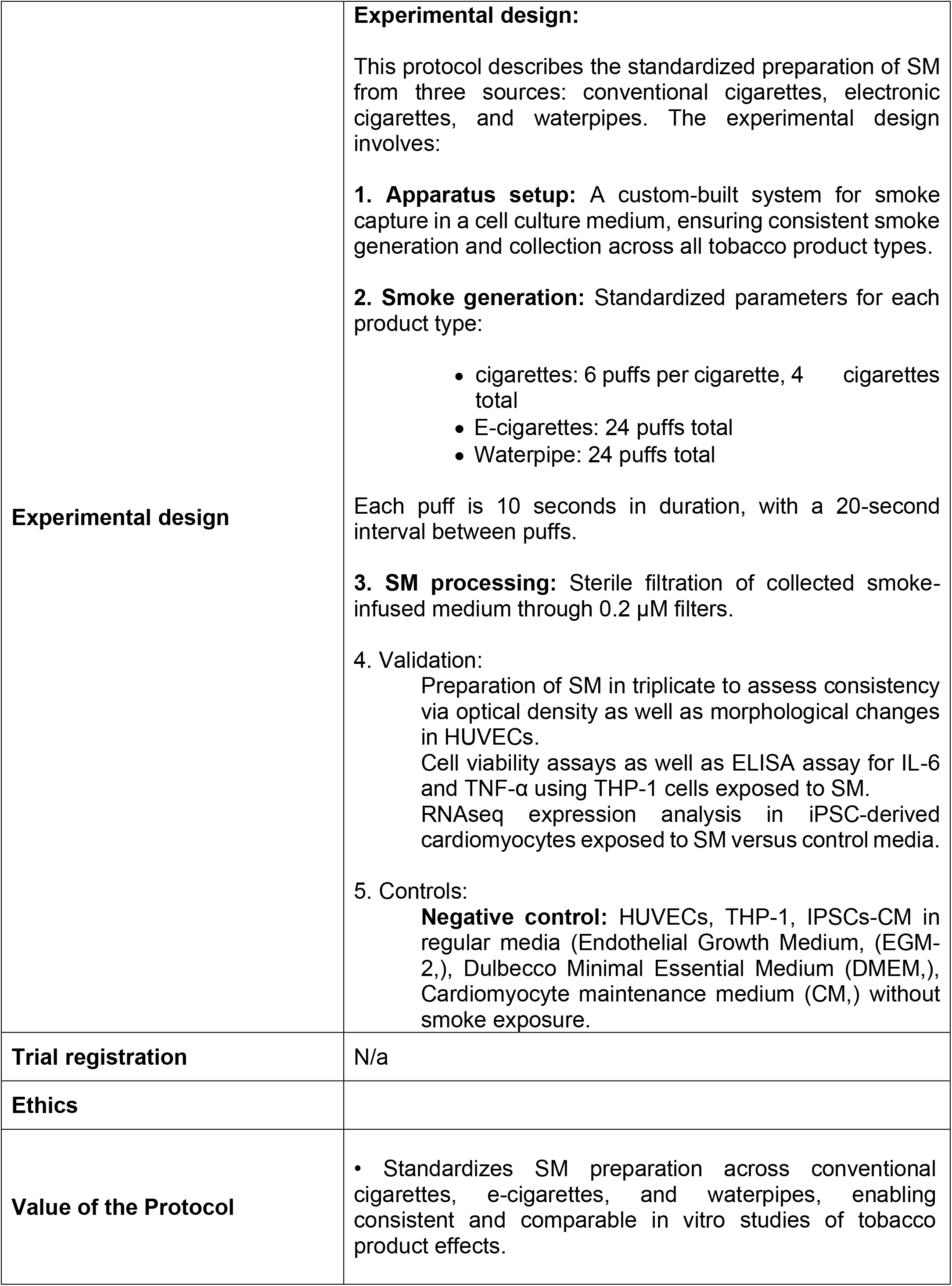

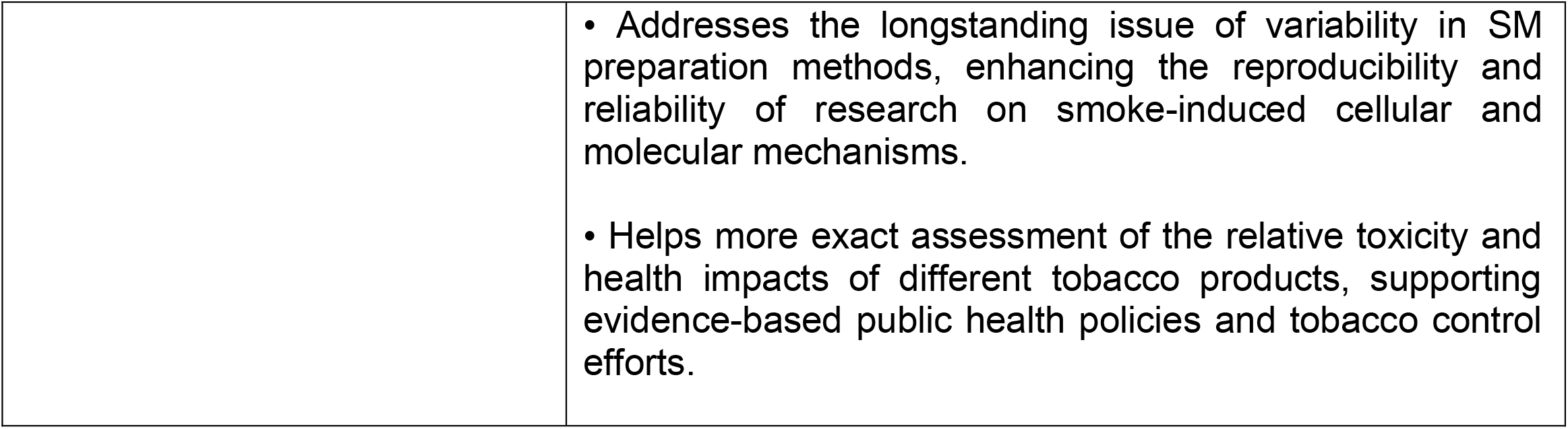

## Background

Tobacco smoking is still a significant global health concern, contributing substantially to morbidity and mortality, particularly concerning cardiovascular and respiratory diseases Roth et al., 2019). Extensive research has documented the detrimental effects of cigarette smoke on human health, implicating it in the pathogenesis of many conditions including atherosclerosis, chronic obstructive pulmonary disease, lung cancer, and exacerbations of asthma **(Bhat et al., 2020; Baneras et al.,2022)**. The complex composition of cigarette smoke, making up over 7,000 chemical compounds including reactive oxygen species, nicotine, and various particulates, underlies its involvement in critical pathological processes such as oxidative stress, inflammation, endothelial dysfunction, and cardiomyocyte injury **(Sun et al., 2016; Kaur et al., 2018)**. In recent years, alternative tobacco products such as electronic cigarettes (e-cigarettes) and waterpipes have gained popularity, particularly among younger populations. While often perceived as less harmful than conventional cigarettes, emerging evidence suggests that these products also pose significant health risks. E-cigarettes have been associated with acute lung injury, cardiovascular dysfunction, and potential long-term respiratory effects **(Gotts et al., 2019; Münzel et al., 2020)**. Similarly, waterpipe smoking has been linked to increased risk of respiratory illnesses, cardiovascular diseases, and certain cancers **(Waziry et al., 2016)**. In vitro models using SM have appeared as invaluable tools for investigating the cellular and molecular mechanisms underlying smoking-related diseases. These models enable researchers to expose relevant cell types, including endothelial cells, airway epithelial cells, and immune cells, to SM derived from various sources such as CSE, ECS, and WPV. This approach eases the elucidation of complex signalling pathways, gene expression alterations, and functional changes induced by smoke components **(Chen et al., 2017; Lee et al., 2018)**. Moreover, these in vitro systems provide a platform for evaluating potential therapeutic interventions and finding novel drug targets for the treatment of smoke-related diseases **(Wang et al., 2020; Agarwal et al., 2021)**. Despite the importance of SM-based research, a significant challenge in this field is the lack of standardized protocols for SM preparation. This inconsistency leads to variability in experimental outcomes and hinders the comparison of data across studies **(Taylor et al., 2020; Cobb et al.,2021)**. Factors such as product type, smoking parameters, and extraction methods can substantially influence the chemical composition and biological activity of the resulting SM, thereby affecting the reproducibility and validity of research findings. To address this critical issue, we propose a comprehensive and optimized protocol for the reproducible generation of SM from multiple sources, including conventional cigarettes, waterpipes (hookahs), and e-cigarettes. This standardized method aims to ensure consistent and reliable outcomes across diverse laboratories and investigations. The protocol provides a detailed approach for creating reproducible in vitro exposure models, which are essential for advancing our understanding of the cellular and molecular effects of various tobacco products on cardiovascular and respiratory health. Implementation of this protocol will enable more correct comparisons of the impacts of different tobacco products, ease investigation of mechanisms underlying smoke-induced pathologies, and support the evaluation of potential therapeutic interventions with greater confidence. By offering a robust foundation for SM generation, this method effectively addresses the pressing need for standardization in smoke-related research. Adoption of this protocol is expected to enhance experimental reproducibility, adopt collaborative research efforts and contribute to the development of effective strategies to mitigate the adverse effects of exposure to various tobacco products.

## Description of protocol

### Materials

Class II Biosafety Cabinet Avidity Science

Chemical Fume Hood: AFA1000, TEL UK

Parafilm M®: Bemis Company, Inc., USA

60 mL Luer-Lok™ Tip Sterile Syringe: Becton Dickinson, USA

250 mL Sterile Polypropylene Conical Centrifuge Tubes: Corning, USA

Tygon® ND 100 Translucent Tubing: Saint-Gobain, USA, 1/4” ID x 3/8” OD

Digital Timer

Aluminium Foil Sheets

Disposable Lighter

Stainless Steel Forceps

50 mL Polypropylene Graduated Beaker:

60 mm Diameter Porcelain Buchner Funnel: Rosen Thaf ATW, German

Marlborough Red Cigarettes: 4 per experiment, filters removed

Al Fakher Molasses Tobacco: Manufacturer, Country, 9.5g per experiment

Instant Lighting Charcoal Discs: Meera Brand, Country, 1 per waterpipe experiment

JUUL Electronic Nicotine Delivery Device: JUUL Labs, USA, fully charged

JUUL Nicotine Pods: Alpine Berry Flavour, 18 mg/mL Nicotine, JUUL Labs, USA

Nalgene Rapid-Flow 0.2 µm Sterile Disposable-Bottom Filter Units: Thermo Scientific, USA

50 mL Sterile Polypropylene Centrifuge Tubes: Corning, USA

Serological Pipettes, Pipette Tips, and Controllers: Eppendorf, Easy Pet 3 and Gilson

All cell culture media, reagents, and materials were obtained from certified sources and prepared using aseptic techniques in a Class II Biosafety Cabinet.

### Apparatus Setup

#### Collection Vessel

Use a 250 mL sterile conical tube for smoke capture in a cell culture medium.

#### Smoke Collection Chamber

Connect a 60 mL syringe via Tygon tubing to the conical tube.

#### Cigarette/E-Cigarette Smoke Collection

Attach a second length of Tygon tubing to the syringe tip to hold the tobacco product in place.

#### Waterpipe Smoke Collection

Connect a 60 mm Buchner funnel via Tygon tubing to the syringe tip to hold the molasses tobacco.

#### Sealing

Seal all tubing connections with parafilm to ensure an airtight system.

#### Negative Pressure Differential

Place a sterile stopper on the top of the conical tube during smoke draws.

The Assembled apparatus should be placed in a chemical fume hood during smoke generation to ensure proper venting. All materials should be handled using aseptic techniques inside a Class II Biosafety Cabinet.

### Smoke Medium (SM) Generation

#### Cigarette Smoke Extract (CSE)

##### Preparation

Use four filtered Marlborough Red cigarettes per replicate. Remove the filters aseptically using forceps.

##### Combustion

Light the end of one cigarette and quickly insert it into the tubing attached to the syringe.

##### Smoke Collection

Pull the syringe plunger back to the 60 mL mark for over 10 seconds to draw the mainstream smoke into the 100 mL cell culture media in the conical tube, standing for one ‘draw’.

##### Repeat Draws

Repeat this process for a total of 6 draws per cigarette, depleting the cigarette.

##### Cycle Continuation

Remove the spent cigarette butt and repeat the process with a new cigarette until all 4 are consumed, totalling 24 draws.

#### Electronic Cigarette (ECS)

##### Device Preparation

Insert a new JUUL nicotine pod (Alpine Berry, 18mg/mL) into a fully charged JUUL device.

##### Mouthpiece Insertion

Gently insert the e-cig mouthpiece into the tubing attached to the syringe.

##### Smoke Collection

Activate the device and draw smoke by pulling the syringe plunger at a rate of 1 draw every 10 seconds for a total of 24 draws.

##### New Replicates

Use a new nicotine pod for each independent replicate.

#### Waterpipe Vapour (WPV)

##### Tobacco Preparation

Weigh and place 9.5g of moist Al Fakher molasses tobacco into the Buchner funnel.

##### Foil Cover

Place a pierced aluminium foil cover over the funnel, creating 100 holes of consistent size with forceps.

##### Charcoal Preparation

Light one charcoal disc using a disposable lighter and place it on the foil cover once fully ashed over.

##### Smoke Collection

Draw smoke using the 60 mL syringe at a rate of 1 draw every 20 seconds for a total of 24 draws.

##### Note

For each experimental condition, the resulting SM holding the total particulate matter from 24 draws is defined as the 100% SM stock. This stock is passed through a 0.2 µm filter unit (Thermo Scientific, USA) using an aseptic technique to remove bacteria and large particulates. The sterile filtered SM is then serially diluted in a complete cell culture medium to the desired test concentrations (e.g., 0.1%, 1%, 10%) based on the experimental design. Diluted SM extracts are prepared from 100% stock for each

#### Experimental Design

##### FDA Guidelines

Follow all applicable FDA guidelines for handling and processing tobacco products and nicotine-containing devices.

##### Human Subject Guidelines

Ensure compliance with ethical standards and guidelines when handling human-derived cell lines or tissues.

#### Data Measurement and Analysis

##### Cell Viability Assays

Assess the impact of SM on cell viability using assays such as MTT or Alamar Blue.

##### Gene Expression Analysis

Employ RNA-seq to analyse changes in gene expression.

##### Protein Analysis

Use ELISA to measure protein levels and modifications.

##### Imaging

Use fluorescence microscopy to visualize cellular changes.

#### Additional Observations and Tips

##### Ensure Airtight Connections

Make sure all connections in the apparatus are airtight to prevent loss of smoke.

##### Aseptic Handling

Perform all steps involving live cells and SM under aseptic conditions to avoid contamination.

##### Proper Venting

Conduct smoke generation steps in a chemical fume hood to ensure proper venting and safety.

##### Personal Protective Equipment

Handle nicotine-containing products with care, using personal protective equipment

## Protocol validation

### Assessing the quality of SM through different modalities and addressing the toxicological effects of the SM on the endothelial cells, i.e., HUVECs

Representative micrographs in Figures **4A** and **4B** illustrate significant changes in the composition and appearance of the medium following exposure to smoke. The SM showed noticeable colour alterations and variations in optical density (O.D.), which were assessed by analysing the absorption spectrum. The normal medium, not exposed to smoke, showed comparatively lower O.D. values. In contrast, the test groups, particularly the WPV group, displayed significantly higher O.D. values, showing a substantial impact from smoke constituents on the medium. To assess the potentially toxic effects of SM, HUVECs were treated for 24 hours as shown in **Figure 4C**. Later imaging at 40x magnification revealed pronounced morphological changes in the HUVECs within the CSE, ECS, and WPV. These cells exhibited CSE loss of normal morphology, with many undergoing shrinkage and detachment, suggesting cytotoxic effects. These observations served as a preliminary benchmark for further detailed investigations. Later results offered comprehensive insights into the arrested cell viability and the regulation of pro-inflammatory cytokines. Additionally, toxicity was assessed at the RNA level, focusing on the expression of genes responsible for cellular damage. These findings underscore the detrimental effects of smoke constituents on endothelial cell health and function.

### Assessing the THP-1 cell viability at 24 hours post SM treatment

Figure 5. stands for a graph in which post-smoking THP-1 cells at 24 hours were analysed to figure out the proliferation of the cells by analysing the % viability of the cells. In this, the Negative control showed the highest cell viability while comparing it to the Positive control as well as different test groups at certain defined concentrations. Moreover, there was a striking reduction in the viability of the THP-1 cells when treated with CSE, especially at higher concentrations which provides the proof-of-concept in line with the altered morphology in the HUVECs as shown in Figure 1C. Furthermore, upon figuring out the proliferation of the THP-1 cells in the ECS and WPV group both treatment groups do not cause higher cytotoxicity to the macrophage cells even at 100% of treatment groups.

**Figure 1.**
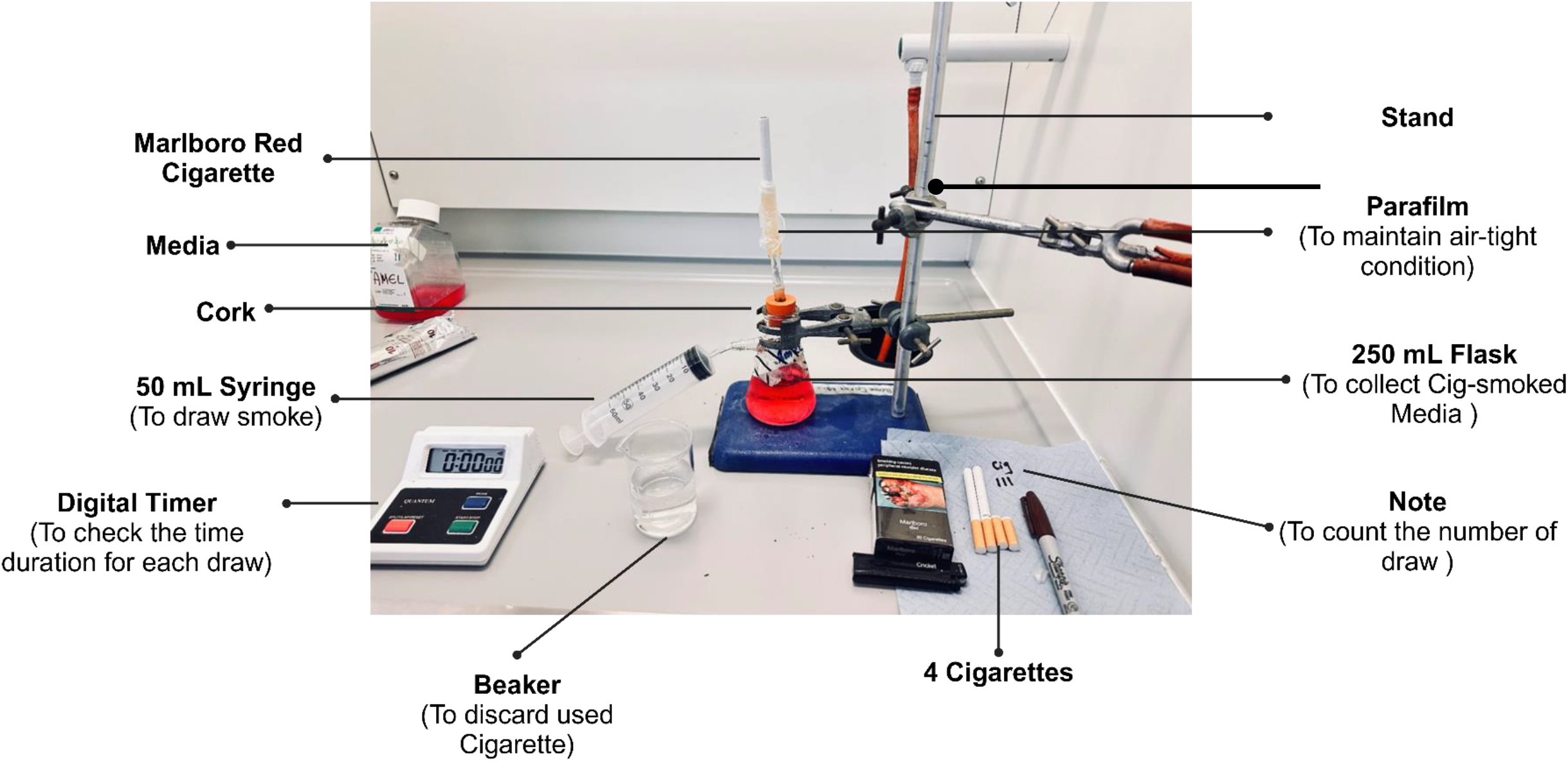
Representative image for the CSE apparatus

**Figure 2.**
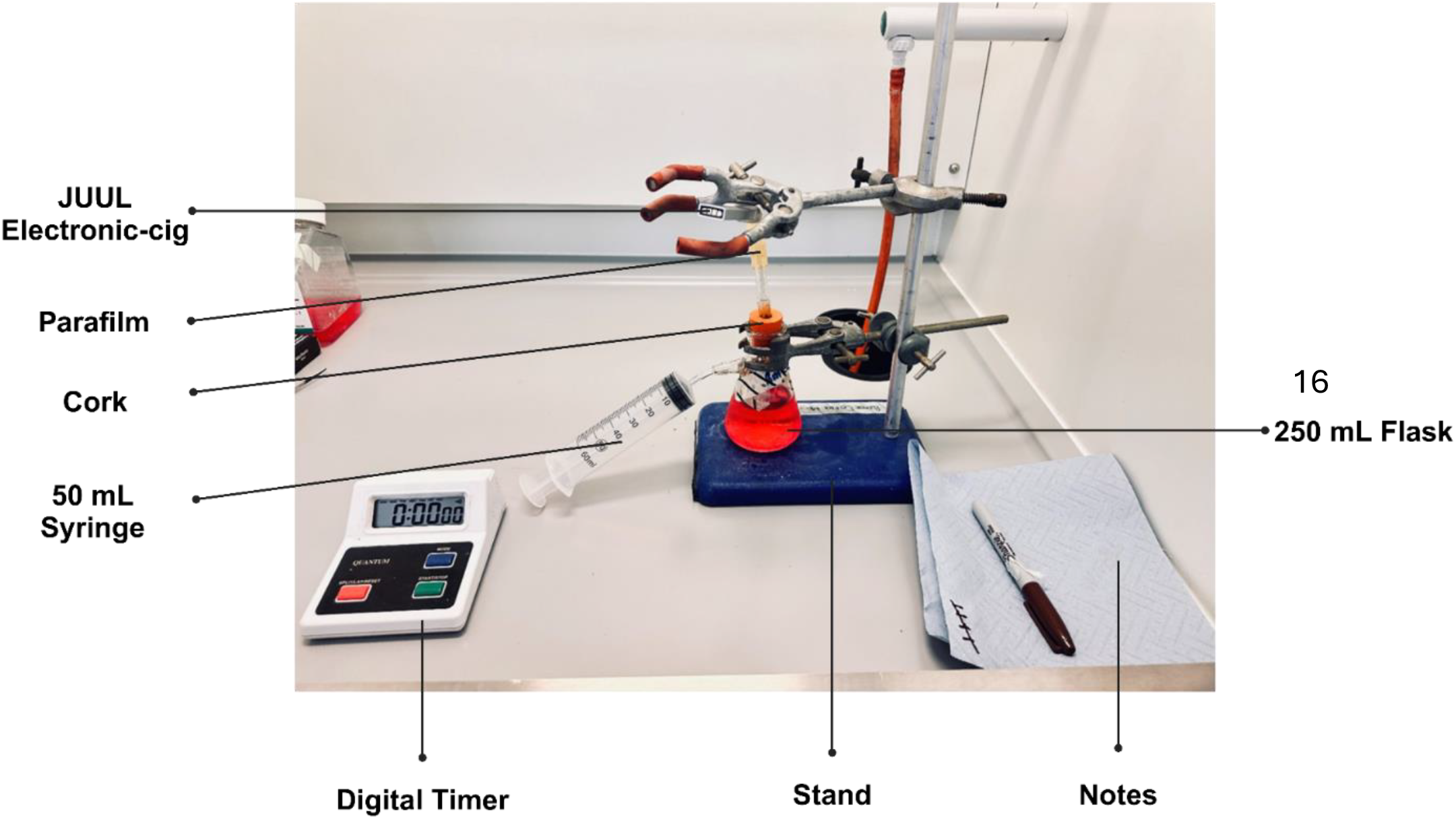
Representative image for the ECS apparatus

**Figure 3.**
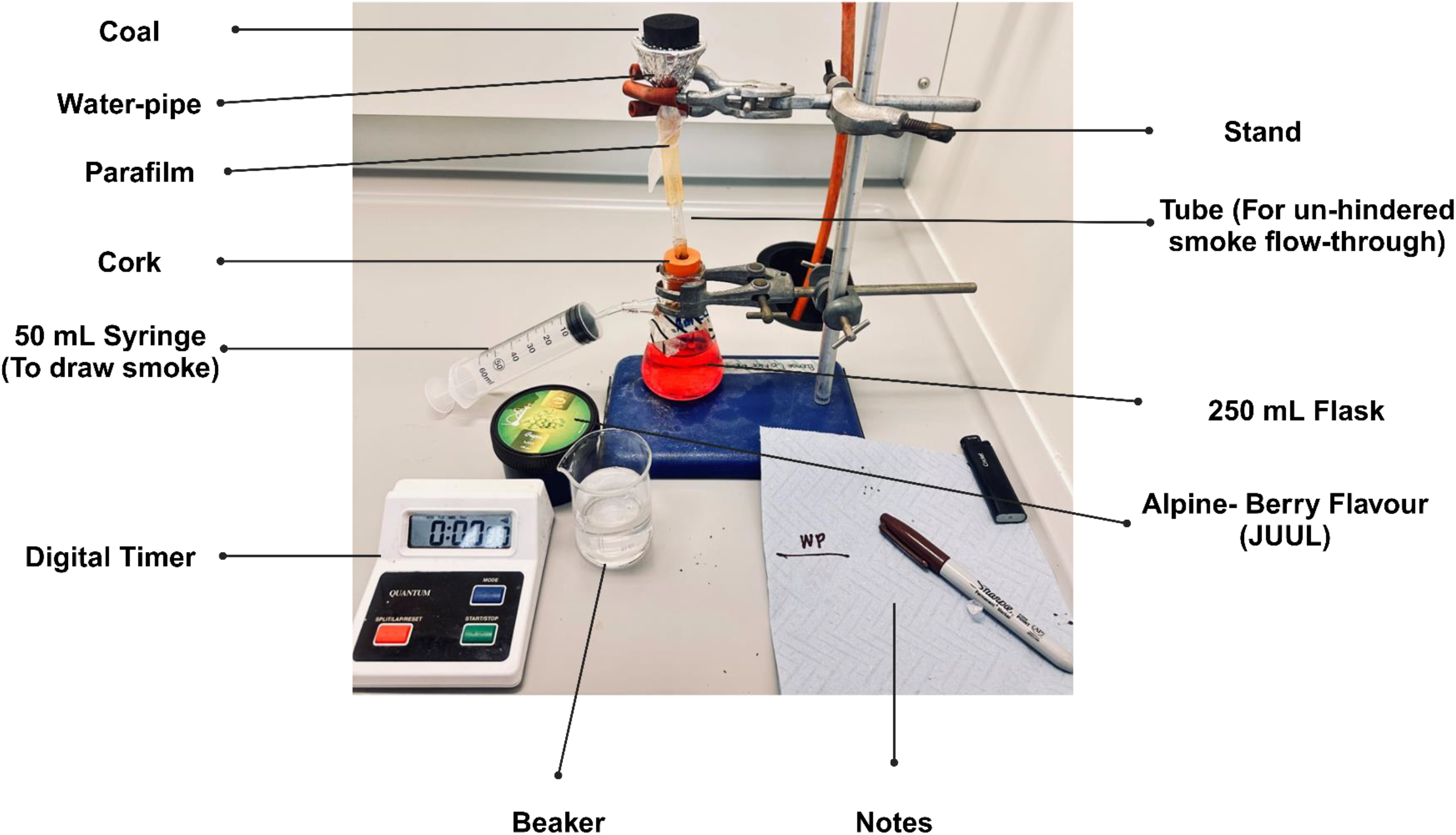
Representative image for the WPV apparatus

### Differential expression of IL-6 Production in THP-1 cells by different SM modalities: Implications for Inflammatory Responses

**Figure 4.**
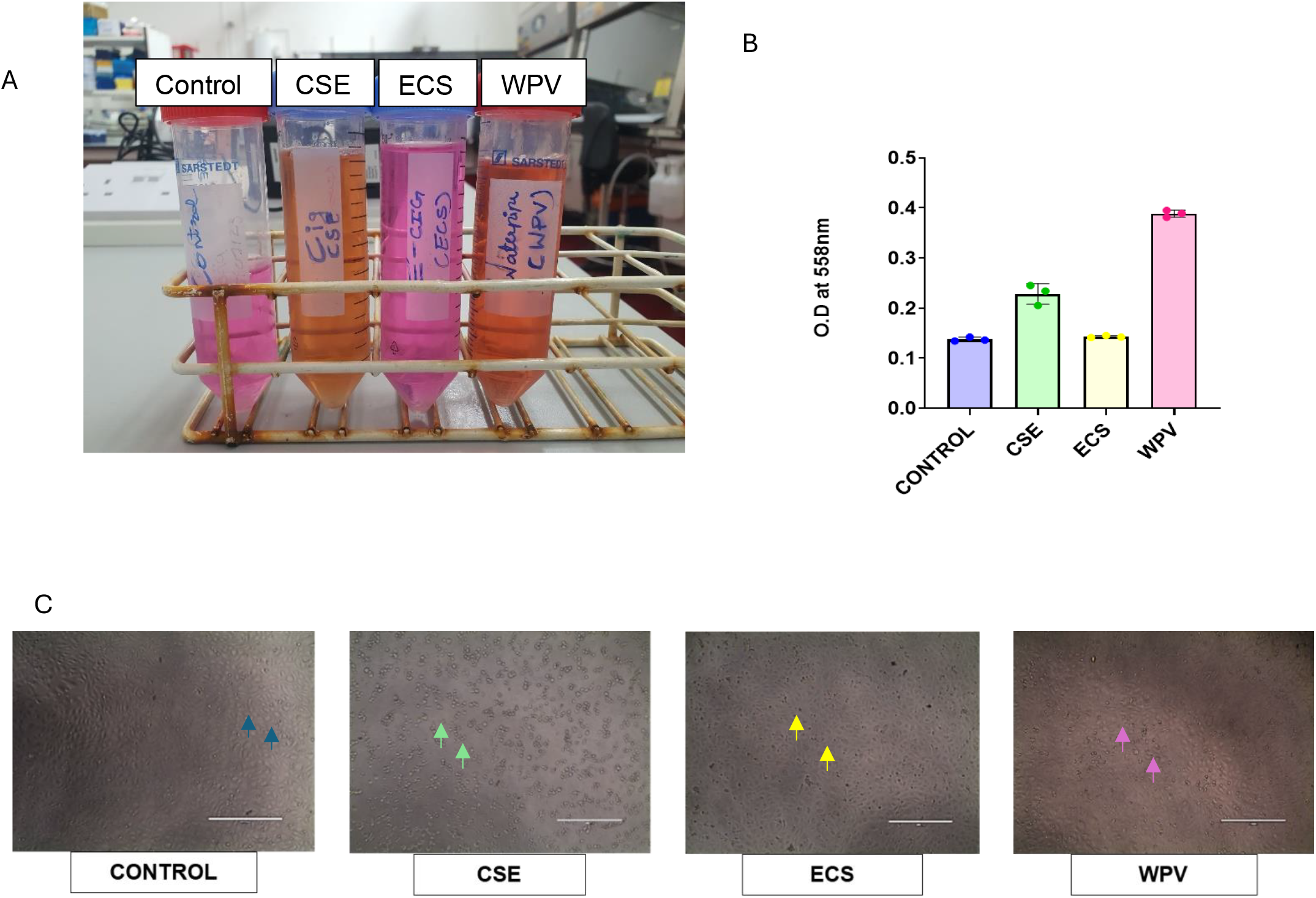
Representative micrographs illustrating alterations in the media following exposure to SM (a), alongside changes in optical density measurements of SM (b) and morphology induced changes by the SM on HUVECs with control media (blue arrows), *CSE*; cigarette smoke extract (green arrows), *ECS*; e-cigarette smoke (yellow arrows) and *WPV*; waterpipe vapor (pink arrows), n=3, Scale bar: 400µM.

**Figure 5.**
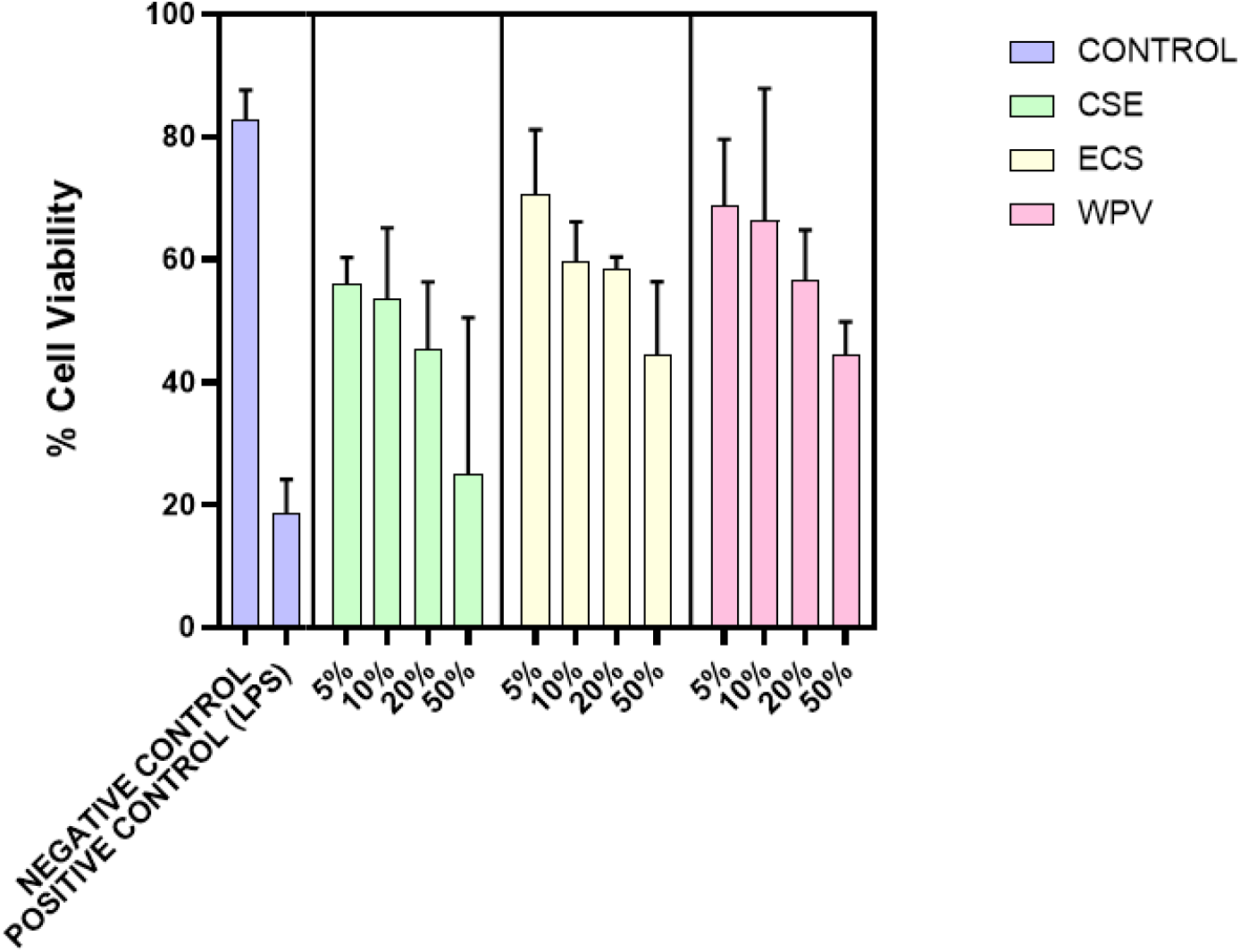
Representative graphs illustrating the Viability of THP-1 cells at the time point of 24 hours when stimulated with Control, *CSE*; cigarette smoke extract, *ECS*; e-cigarette smoke and *WPV*; waterpipe vapor at different concentrations, n=3.

## Results

Figure 6. illustrates the up-regulation of interleukin-6 (IL-6), a potent pro-inflammatory cytokine, in response to stimulation by CSE, ECS, and WPV. The graph shows that WPV induces the highest level of IL-6 production at a 15% concentration, which aligns closely with the ECS data. Both WPV and ECS elicit substantial IL-6 production, with levels reaching approximately 70-80 μg/mL. Interestingly, the CSE group shows unexpectedly low IL-6 production across all smoking modalities tested. This observation leads to the hypothesis that CSE may induce higher levels of cytotoxicity in macrophage cells while simultaneously suppressing IL-6 production. Further investigation into this phenomenon is called for to elucidate the underlying mechanisms. As a positive control, lipopolysaccharide (LPS), a known potent stimulator of IL-6, was used. The LPS group proved a marked up-regulation of IL-6 production when presented to macrophage cells. This result not only confirms the accuracy of the smoking protocol but also provides robust evidence for its reproducibility across various in vitro models. These findings collectively suggest differential effects of various smoking modalities on IL-6 production in macrophage cells, with WPV and ECS showing the most pronounced pro-inflammatory responses. The unexpected results seen with CSE highlight the complexity of cellular responses to different tobacco and nicotine delivery systems, emphasizing the need for further research to fully understand their impact on inflammatory processes.

### RNAseq profiling of iPSC-cardiomyocytes exposed to different SM modalities: Identification of Potential Genes Implicated in Cardiovascular Disease

**Figure 6.**
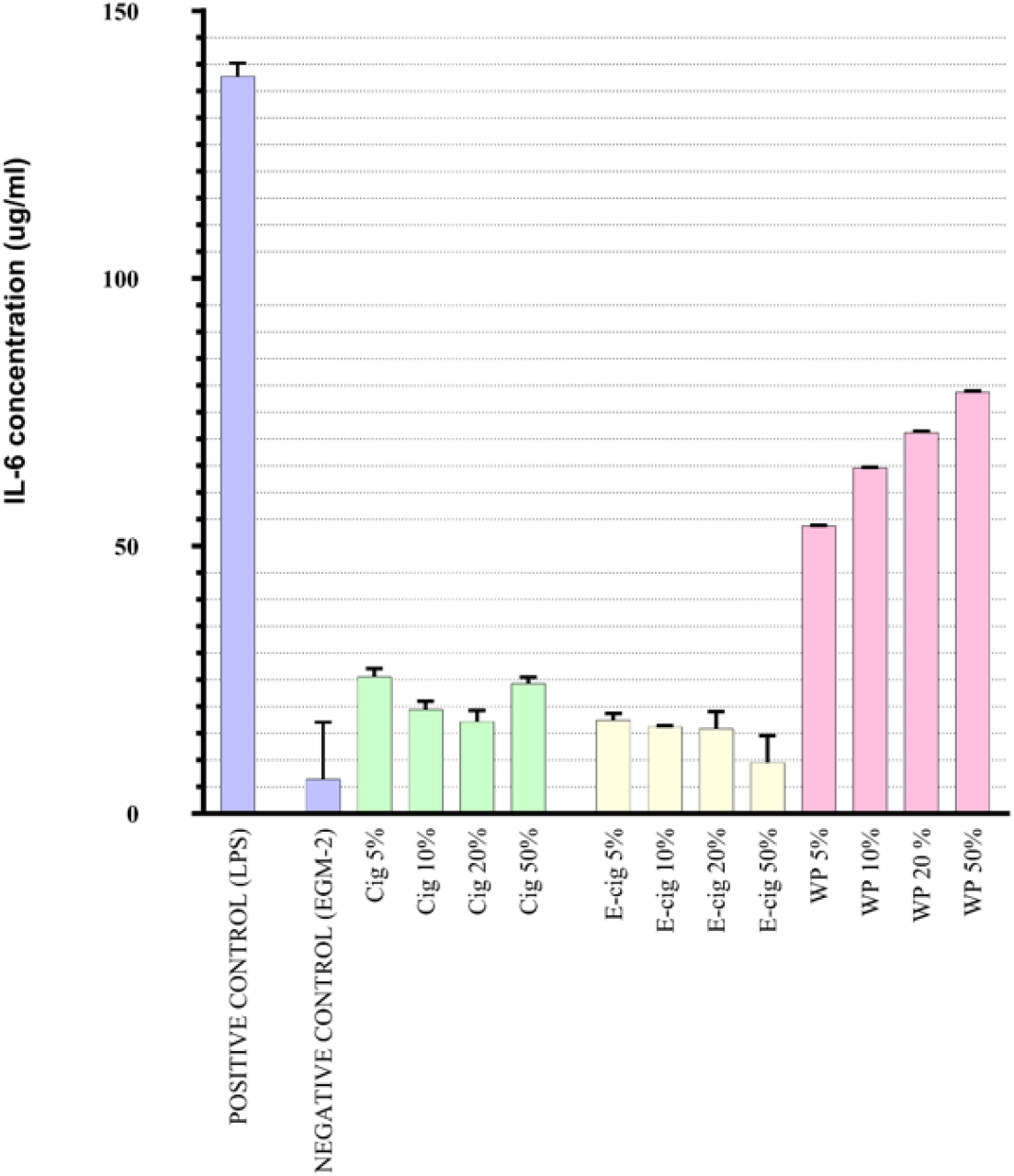
Representative graphs illustrating the proliferation of IL-6 at the time point of 24 hours when stimulated with Control, CSE, ECS and WPV at certain different concentration. n=3

## Results

**Figure 7A** shows RNAseq analysis of iPSC-CMs exposed to CSE, ECS, and WPV, which revealed significant differential gene expression changes compared to controls. CSE exposure induced the most pronounced changes, with 1,570 upregulated and 7,710 downregulated genes. ECS led to more modest alterations (484 upregulated, 569), while WPV resulted in 2,048 upregulated and 2,988 downregulated genes. Volcano plots visually stood for these expression changes, clearly delineating significantly altered genes from those unchanged across smoke types.

**Figure 7a,7b.**
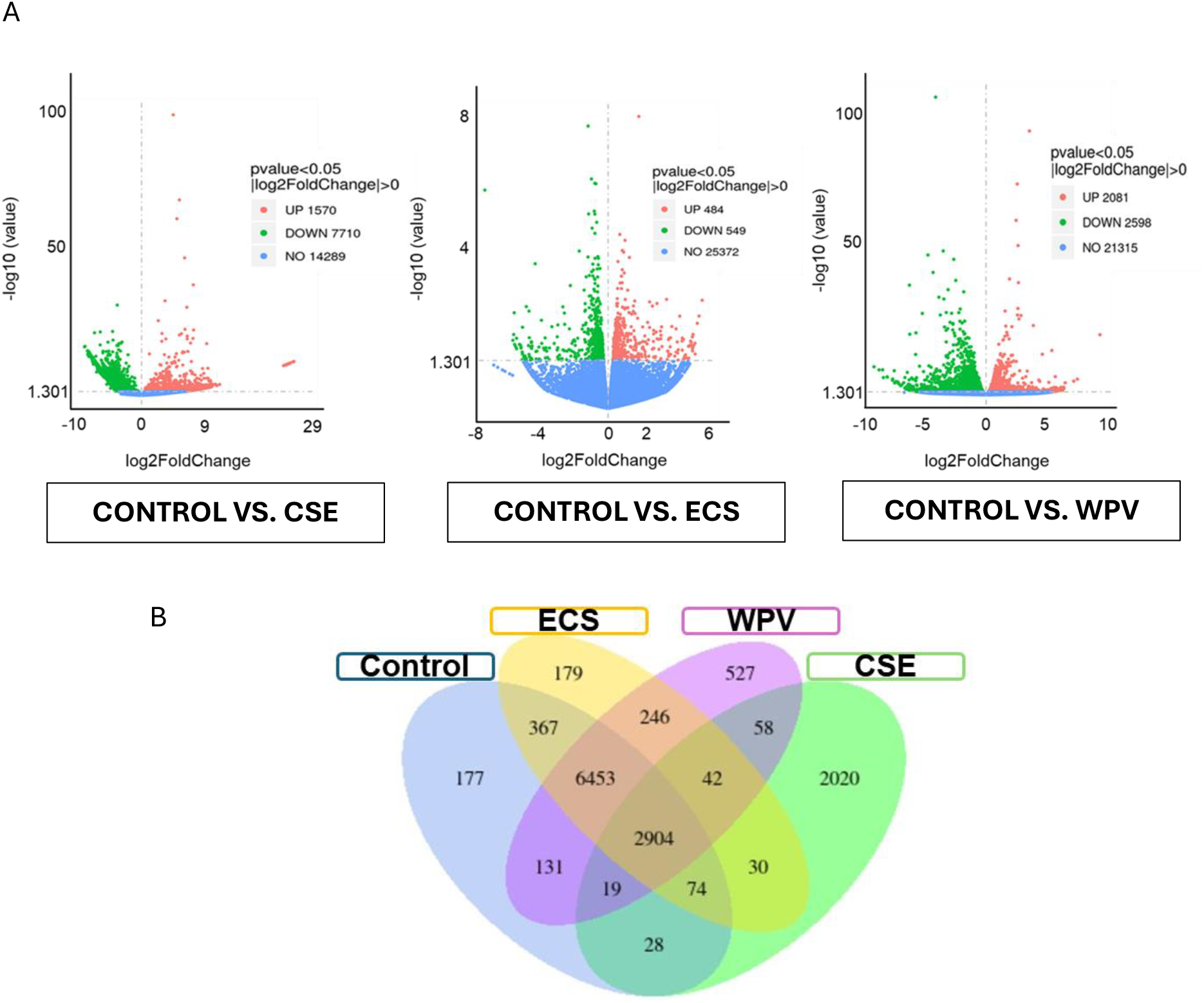
Volcano Plot of Differential Gene Expression Between Control vs. cigarette smoke extract CSE), E-cigarette smoke (ECS) and waterpipe vapour (WP) on iPSC-cardiomyocytes. The Venn diagram ustrates the overlap of differentially expressed genes in iPSC-cardiomyocytes exposed to CSE, ECS, and PV. Each section represents the unique and common genes among the three groups, highlighting potential enes implicated in the development of CVDs due to smoke exposure.

Venn diagram analysis highlighted unique and shared differentially expressed genes among the three smoke exposure groups, providing insights into potential common pathways implicated in smoke-induced cardiovascular damage **Figure 7B**. The substantial transcriptomic changes seen across all smoke types underscore the broad impact of various smoke exposures on iPSC-CM gene expression. These findings establish a foundation for further investigation into the molecular mechanisms underlying smoke-induced cardiovascular diseases especially 42 genes which are shared amongst each SM group and absent in the control group are of the main research interest in future work while demonstrating the efficacy of the employed SM protocol in consistently inducing significant transcriptomic alterations in iPSC-CMs across different SMs (Smoke Medium).

### Limitations

While this protocol offers a standardized approach for generating smoke extract media from various tobacco products, several limitations should be considered. The inherent variability in tobacco product composition may affect reproducibility across studies. The fixed smoking topography simulation may not accurately be all real-world scenarios, potentially limiting translational relevance. Filtration steps could lead to the loss of volatile compounds, altering the extract’s composition. The protocol does not address potential diffusing effects on the extract or temperature considerations during generation, which may affect chemical composition. Despite standardization efforts, inter-operator variability is still a potential issue. Additionally, the in vitro nature of the exposure model may not fully capture the complexities of in vivo smoke exposure. The protocol’s optimization for specific product types may require adaptation and validation for significantly distinctive designs or formulations. These limitations should be carefully considered when interpreting results and comparing findings across studies, and further refinement may be necessary to enhance the method’s robustness.

## CRediT author statement

**Amel-Al Hashimi**: Conceptualization, Methodology, Validity, Data Curation, Software. **Jagrut Shah**: Validity tests, Data curation, Writing-Original draft preparation. **Dr. Mohammed Meah**: Visualization, Investigation. Dr **Roger Carpenter**: Supervision., **Prof. Winston Morgan**: Supervision. **Dr. Prashant J. Ruchaya**: Conceptualization, Methodology, Reviewing and Editing.

## References

1. Agarwal, Amit R., et al. “Short-Term Cigarette Smoke Exposure Leads to Metabolic Alterations in Lung Alveolar Cells.” American Journal of Respiratory Cell and Molecular Biology, Mar. 2014, p. 140313133749009, 10.1165/rcmb.2013-0523oc. Accessed 6 Jan. 2021.

2. BAÑERAS, Jordi, et al. “Medio Ambiente Y Salud Cardiovascular: Causas, Consecuencias Y Oportunidades En Prevención Y Tratamiento.” Revista Espanola de Cardiologia (English Ed.), vol. 75, no. 12, Dec. 2022, pp. 1050–58, 10.1016/j.rec.2022.05.030. Accessed 19 Sept. 2023.

3. Bhat, Tariq A., et al. “An Animal Model of Inhaled Vitamin E Acetate and EVALI-like Lung Injury.” New England Journal of Medicine, Feb. 2020, 10.1056/nejmc2000231. Accessed 3 Mar. 2020.

4. Chen, H. S., et al. “Maternal E-Cigarette Exposure in Mice Alters DNA Methylation and Lung Cytokine Expression in Offspring.” American Journal of Respiratory Cell and Molecular Biology, vol. 58, no. 3, American Thoracic Society, Sept. 2017, pp. 366–77, 10.1165/rcmb.2017-0206rc.

5. Cobb, Caroline O., et al. “Effect of an Electronic Nicotine Delivery System with 0, 8, or 36 Mg/ML Liquid Nicotine versus a Cigarette Substitute on Tobacco-Related Toxicant Exposure: A Four-Arm, Parallel-Group, Randomised, Controlled Trial.” The Lancet Respiratory Medicine, vol. 9, no. 8, Aug. 2021, pp. 840–50, 10.1016/s2213-2600(21)00022-9.

6. Gotts, Jeffrey E., et al. “What Are the Respiratory Effects of E-Cigarettes?” BMJ, vol. 366, no. 5275, Sept. 2019, p. l5275, 10.1136/bmj.l5275.

7. Kaur, Gurjot, et al. “Mechanisms of Toxicity and Biomarkers of Flavouring and Flavour Enhancing Chemicals in Emerging Tobacco and Non-Tobacco Products.” Toxicology Letters, vol. 288, 2018, pp. 143–55, 10.1016/j.toxlet.2018.02.025.

8. Lee, Hyun-Wook, et al. “E-Cigarette Smoke Damages DNA and Reduces Repair Activity in Mouse Lung, Heart, and Bladder as well as in Human Lung and Bladder Cells.” Proceedings of the National Academy of Sciences, vol. 115, no. 7, Jan. 2018, pp. E1560–69, 10.1073/pnas.1718185115.

9. Münzel, Thomas, et al. “Effects of Tobacco Cigarettes, E-Cigarettes, and Waterpipe Smoking on Endothelial Function and Clinical Outcomes.” European Heart Journal, vol. 41, no. 41, June 2020, 10.1093/eurheartj/ehaa460.

10. Roth, Gregory A., et al. “Global Burden of Cardiovascular Diseases and Risk Factors, 1990-2019: Update from the GBD 2019 Study.” Journal of the American College of Cardiology, vol. 76, no. 25, Dec. 2020, pp. 2982–3021, 10.1016/j.jacc.2020.11.010.

11. Sun, Desheng, et al. “Cigarette Smoke-Induced Chronic Obstructive Pulmonary Disease Is Attenuated by CCL20-Blocker: A Rat Model.” Croatian Medical Journal, vol. 57, no. 4, Aug. 2016, pp. 363–70, 10.3325/cmj.2016.57.363. Accessed 7 Jan. 2020.

12. Taylor, Mark, et al. “E-Cigarette Aerosols Induce Lower Oxidative Stress In Vitro When Compared to Tobacco Smoke.” Toxicology Mechanisms and Methods, vol. 26, no. 6, July 2016, pp. 465–76, 10.1080/15376516.2016.1222473. Accessed 26 Jan. 2022.

13. Wang, Qixin, et al. “E-Cigarette-Induced Pulmonary Inflammation and Dysregulated Repair Are Mediated by NAChR α7 Receptor: Role of NAChR α7 in SARS-CoV-2 Covid-19 ACE2 Receptor Regulation.” Respiratory Research, vol. 21, no. 1, June 2020, 10.1186/s12931-020-01396-y. Accessed 26 Aug. 2020.

14. Waziry, Reem, et al. “The Effects of Waterpipe Tobacco Smoking on Health Outcomes: An Updated Systematic Review and Meta-Analysis: Table 1.” International Journal of Epidemiology, Apr. 2016, p. dyw021, 10.1093/ije/dyw021.

